# A mechanistic spatiotemporal model for drug resistant infections

**DOI:** 10.1101/2022.03.03.482844

**Authors:** Tamsin E. Lee, Pablo M. De Salazar

**Affiliations:** Swiss Tropical and Public Health Institute, Basel, Switzerland; University of Basel, Basel, Switzerland

## Abstract

Effective public health strategies for preventing infectious diseases require a deep understanding of transmission dynamics. However, epidemiological data alone is often insufficient for guiding public health action, as transmission patterns emerge from complex human-environment interactions. Additionally, collecting empirical data is frequently limited by cost and logistical challenges, especially in low-resource settings. Consequently, health policy increasingly relies on inference models that account for spatial and temporal dimensions, confounding factors, and uncertainty.

In this work, we introduce a novel hierarchical mechanistic Bayesian model based on ecological diffusion to improve epidemiological inference for spreading pathogens. This model is specifically designed to reveal underlying spatiotemporal dynamics from empirical data, explicitly accounting for sampling bias and uncertainty. To demonstrate its effectiveness, we apply the model to simulated data of a drug-resistant pathogen and formally compare its performance to state-of-the-art epidemiological regression models.

Our findings show that the model accurately identifies transmission patterns and is particularly effective at pinpointing transmission hotspots and predicting spread pathways—critical capabilities for controlling drug-resistant pathogens. While reliable monitoring remains challenging in low-resource environments, this model addresses demographic sampling bias and spatiotemporal complexities, though concentrated sampling in presumed high-risk areas can sometimes distort transmission insights.

The proposed framework can inform targeted public health interventions in the early stages of resistance emergence, offering valuable guidance for resource-constrained settings.

## Introduction

Public health strategies aiming to prevent and control communicable diseases need primarily to identify the emergence hotspots and predict their pathways of spread ***Brookes et al. (2015)***. Via epidemiological surveillance, health-systems can track the endemic transmission patterns as well as outbreaks and epidemics of emerging pathogens ***World Health Organization (2023)***. However, for monitoring to lead to effective action, it requires a sufficient number of high-quality empirical observations, ensuring that any inferences made are reliable. This is particularly challenging in emerging and reemerging pathogens ***Roberts (2019)***, as well as in settings where surveillance systems are not sufficiently resourced ***Iskandar et al. (2021)***. A particularly relevant case is the spread of drug-resistant pathogens, because of the global threat and the potential severe consequences that they pose to human health. Once drug-resistant pathogens establish in a population, our ability to manage the resulting infections and to mitigate their spread can be strongly reduced ***Boni et al. (2008)***. At early stages, resistant-clones typically emerge in hotspots, such as health centers or transportation hubs, to further spread into the population. Rapidly identifying these resistance hotspots, and the underlying transmission patterns over the population is a critical first step to mitigate the impact of resistance.

Transmission dynamics in drug-resistant infections are still not fully understood. For instance, drug-sensitive and drug-resistant infections compete for hosts—a process influenced by factors such as healthcare access and socioeconomic conditions, including limited availability of high-quality medications and poor adherence to treatment protocols, particularly in low-resource settings. Dynamic epidemiological models can offer crucial insights into these complexities; however, to be truly effective, they must explicitly address confounding factors such as sampling bias, incorporate spatial and temporal dimensions that capture transmission patterns, and account for inherent uncertainties ***Knight et al. (2019)***. Since the early 2000s, hierarchical mechanistic models have been developed for predicting the spread of ecological process ***Wikle (2003)***; ***Wikle and Hooten (2006)***. Based on careful assumptions which influence the spatiotemporal dynamics and data collection process, each model is tailored to the specific problem and dataset, making them sophisticated and powerful models ***Conn et al. (2015)***.

In the era of digital platforms and novel technologies, classical surveillance systems can incorporate new sources of information that improve our ability to predict the transmission, the distribution and the disease burden of infectious diseases, such as human mobility networks ***Buckee et al. (2013)***, molecular surveillance with pathogen’s evolutionary components ***Gardy and Loman (2018)***, or seroprevalence analysis.

We introduce a novel hierarchical mechanistic Bayesian model based on ecological diffusion ***Hefley et al. (2017)***, marking its first application in antimicrobial resistance epidemiology. This model is designed to uncover the underlying spatiotemporal dynamics from empirical data while explicitly accounting for demographic sampling bias (e.g., age) and uncertainty. This approach allows for the quantification of hotspot contributions to the spread of resistant pathogens within populations. To demonstrate its effectiveness, we evaluate the model using simulated data and compare its performance against state-of-the-art generalised additive models with spatial and temporal components. Furthermore, we show that epidemiological sampling bias—where surveillance efforts are concentrated in regions anticipated to harbor resistant pathogens—distorts the inferred spatiotemporal dynamics in both modeling approaches. The proposed framework is broadly applicable to any drug-resistant pathogen with available spatiotemporal occurrence data, including molecular markers, genome sequencing ***Mari et al. (2021)***, or phenotypic studies ***Davison et al. (2000)***.

## Results

We generated ten datasets by sampling different percentages (5%, 10%, and so on) of the region, both with and without epidemiological bias. Here, ‘biased’ sampling is defined as selecting individuals more likely to test positive for the condition. Using these datasets, the model estimates key parameters, *β* and ***α***, to predict the density of positive cases, which varies by location and over time.

To evaluate the model performance, we calculate the root mean square error (RMSE), which represents the difference between the predicted and actual number of positive cases. We analyse this error over a 20 year period to assess how accurately the model predicts case density across both space and time (see Fig. 6). Additionally, we examine the model’s accuracy in recovering key parameters, *β* and ***α***.

The model has three levels. The first level accounts for various covariates, such as age, that might affect the likelihood of an individual being selected for sampling. This influence is represented by the parameter *β*. Under epidemiological biased sampling, where individuals likely to test positive are prioritised, the model struggles to accurately recover *β*, limiting its ability to provide insights into factors like age and their impact on sampling probability. However, with unbiased sampling of just 5% of the region, the model can accurately estimate how these covariates affect selection probability, successfully recovering *β*.

For each of the ten datasets, we apply two models: a hierarchical Bayesian model (HBM) and a generalised additive model (GAM). Both models predict the spatial distribution of positive cases over the 20 year period, starting by estimating key parameters. We assess model accuracy in two ways. First, we calculate the RMSE to determine how much the predicted density of positive cases deviates from the true density across the entire period (see Fig. 6). A low RMSE indicates that all estimated parameters closely approximate their true values.

Under unbiased sampling, the HBM achieves a low RMSE, performing well with only 5% of the region sampled. Increasing the sample size beyond 5% adds little to accuracy. However, with epidemiological biased sampling, where individuals likely to test positive are prioritised, even a 30% sample yields inaccurate, misleading results. When this biased sampling covers only 5% of the region, the model fails to converge, raising concerns about its reliability in biased sampling conditions for spatiotemporal predictions.

In contrast, the GAM, which explicitly accounts for spatial and temporal changes (using *η*_*s*_ and *η*_*t*_ in Eq. 8), shows consistently high RMSE values that do not improve with increased sample size. In fact, the GAM’s RMSE is higher than the HBM’s across all sample sizes, indicating that the HBM - using just 5% unbiased sampling—produces more accurate estimates than the GAM, even when the latter uses an unbiased 30% sample.

We further assess the models by examining their accuracy in recovering key parameters. Accurate ***β*** estimates mean the model has effectively captured how factors like age influence sampling probability, while precise ***α*** and ***γ*** estimates reflect the model’s ability to capture the local and neighboring spread of infections. Additionally, accurate ***θ*** and ***ϕ*** estimates indicate the model’s success in identifying specific hotspots (areas with high case concentrations).

The parameter *θ*_*i*_ (*i* = 1, 2, 3, 4, 5) represents the hotspot’s intensity, or the number of cases originating from that area, while *ϕ*_*i*_ describes the spatial spread of infections around the hotspot. Together, these parameters reveal spatial dispersion patterns, independent of external factors like transportation. For instance, hotspot 1 had a large *θ*_1_ (many cases) but a small *ϕ*_1_ (limited spread), suggesting a remote healthcare facility with limited accessibility and care quality.

With unbiased sampling, the HBM accurately recovers all parameters, especially *β*, representing the influence of individual covariates like age. This accuracy aligns with expectations, as unbiased sampling avoids distortion and results in a low overall RMSE.

In contrast, when applying the GAM under unbiased sampling, *β* is accurately recovered at all sample sizes because the GAM directly models sampling probability based on individual characteristics (Eqns. 7 and 8). However, with epidemiological biased sampling, the accuracy of *β* recovery diminishes and depends on the sample proportion. In these cases, the GAM performs better than the HBM at recovering *β* but at the expense of higher overall RMSE.

## Discussion

Effectively combating drug-resistant infections would require epidemiological modelling frameworks that can capture the underlying dynamics driving the spread of resistance, especially in relation to resistance hotspots like healthcare centers or transport hubs. Relying solely on traditional monitoring tends to prioritise high-risk regions, potentially overlooking emerging areas where resistance could also be taking hold. Hierarchical mechanistic models, which have been successfully used in ecological studies to track the spread of species or diseases, present an underutilized yet valuable approach in epidemiology. In this study, we adapted a mechanistic model originally designed for species spread to map the transmission of drug-resistant pathogens in a human population. This model, structured across three levels, offers a comprehensive framework for understanding and predicting resistance spread patterns in a defined region.

The first level accounts for individual demographic factors (such as age) that influence a patient’s probability of being sampled, addressing potential demographic biases in sample selection. The second level captures the spatiotemporal dynamics of infection density through a partial dif-ferential equation (PDE) that combines diffusion between regions with local growth dynamics. Enhanced by an additional term to account for resistance emergence from specific hotspots, this level allows for the realistic modelling of disease spread. The third level estimates model parameters and incorporates uncertainty, enabling a more nuanced understanding of resistance dynamics. Testing this model on simulated data demonstrated its ability to accurately recover key parameters, supporting its potential effectiveness.

To illustrate model performance, we simulated data representing a remote region with a single transport hub and four healthcare centers. In this setting, movement is primarily local, reflecting limited road access and short-range transmission. However, this model can be adapted for scenarios where infection transmission spans greater distances. For instance, in a region with an established road network enabling long-distance spread, the current PDE could be modified to use a fractional Laplacian term to represent ‘superdiffusion’ dynamics, enhancing the model’s adaptability to various real-world landscapes. In larger regions with multiple transport hubs, the model could instead incorporate a directed network analysis framework to better represent regional movement patterns and potential spread.

Our hierarchical mechanistic model outperforms regression-based approaches like generalised additive models (GAMs), as it directly incorporates mechanistic components essential to the true dynamics of disease spread. Even advanced models such as logistic Gaussian processes or stacked Gaussian processes lack these explicit mechanistic elements, making them less suited for capturing and interpreting the underlying causes of transmission. In tests with unbiased sampling, our mechanistic model more accurately estimated positive case density than the GAM, emphasising the value of mechanistic elements that align with observed transmission dynamics.

The influence of sampling bias - where sampling is concentrated in high-risk regions - further underscores the importance of mechanistic modelling. With epidemiological biased sampling of only 5% of the region, the model failed to converge, demonstrating how inadequate or highly concentrated sampling can misrepresent infection dynamics. Even with larger biased sample sizes, results did not yield an accurate ranking of resistance hotspots, which could lead public health resources to be misdirected. For example, biased sampling may inaccurately identify a healthcare center as a resistance hotspot due to high patient density or location near other hotspots, rather than reflecting true transmission dynamics. In such cases, resources would be more effectively allocated based on unbiased sampling data, which offers a clearer view of hotspot contributions.

Evolving data collection methods provide new opportunities for refining sampling strategies. Our findings support epidemiological unbiased (random) sampling, especially at early stages, as it offers a more reliable understanding of the spread mechanisms for drug resistance. This approach could ultimately extend the lifespan of existing treatments by informing more accurate and timely interventions. Interestingly, with biased sampling, the GAM achieved higher accuracy than our model but failed to accurately capture dependencies between resistance and prevalence. This reinforces the need for unbiased data, as concentrated sampling in presumed high-risk areas impacts model reliability for spatiotemporal predictions, regardless of the modelling framework used.

Our hierarchical mechanistic model is flexible and applicable beyond drug-resistant infections. Hotspot detection is essential for understanding other infectious diseases, and the model’s structure allows for dynamic adjustments, such as adding or removing hotspots over time by modifying the final term in Eq. 4. It can also estimate hotspot locations by treating them as model parameters, making the model adaptable to evolving data landscapes. This flexibility enables comparisons between disease hotspots and resistance hotspots, which can provide key insights for targeted public health interventions.

## Methods and Materials

We modify the hierarchical mechanistic model from ***Hefley et al. (2017)***, which modelled the spread of chronic wasting disease in white tailed deer in the southwestern portion of Winconsin. This model used presence/absence data, with a single origin hotspot based on where the disease was first detected. It assumed that the hotspot only contributed the disease to the population at the first time interval. In our modified model, we allow for multiple hotspots, which continually contribute drug resistant pathogens into the population. Furthermore we use count data (not presence/absence), as in ***Hooten and Wikle (2008)***; ***Conn et al. (2015)***.

The model uses an aggregate of resistance surveillance studies. For each study, there is the number of infected patients, and from these tested patients, the number who carry drug resistant pathogens (identified by means such as a carrying a mutation). For brevity, we refer to sampled patients who carried drug resistant pathogens as positive patients, and those who did not carry drug resistant pathogens as negative patients.

We assume that each location, **s** = (*s*_1_, *s*_2_), has a weight that depends on its distance from a hotspot. This weight is greater when there is a greater chance that an infected person develops a mutation that infers resistance to treatment. The weights, *ω*(**s**), are defined by

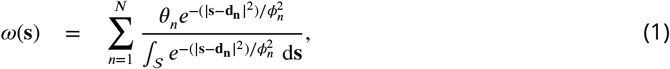

where the magnitude of resistant pathogens contributed by hotspot *n* is *θ*_*n*_, and these resistant pathogens disperse at rate *ϕ*_*n*_ from the hotspot. Eq. 1 is the sum, over *n*, of scaled bivariate Gaussian kernels with compact (truncated) support centred at a point with coordinate **d**_**n**_ where |**s** − **d**_**n**_| is the distance. The units of this distance depends on the size of the region being investigated. An example of *ω*(**s**), where there are five hotspots (*N* = 5) is shown in Fig. 1.

**Figure 1.**
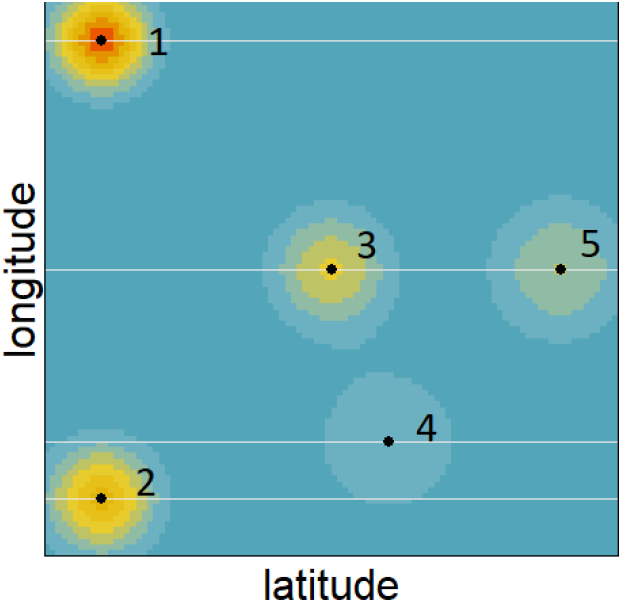
An example of the spatial weightings which are greater (red) where drug-resistant infections are more likely via five hotspots with magnitude *θ*_1_ ≥ *θ*_2_ ≥ *θ*_3_ ≥ *θ*_4_ ≥ *θ*_5_ and dispersal *ϕ*_4_ ≥ *ϕ*_5_ ≥ *ϕ*_3_ ≥ *ϕ*_2_ ≥ *ϕ*_1_ (see the definition of *ω*(**s**) from Eq. 1).

We now provide details of the hierarchical mechanistic model which has separate components to capture uncertainty in data collection, the spatiotemporal transmission dynamics, and uncertainty in the parameters. Following this, we provide details of the numerical implementation, beginning with the algorithm. Then, to thoroughly demonstrate our model, we simulate data over a region that is split into 10,000 grid points (a square that is 100 by 100).

For comparison, we applied a generalised additive model (GAM) to the same simulated data. Generalised additive models are a well-developed and sophisticated statistical tool, where spatial and temporal components can be explicitly included, and the effect of covariates are readily quantified. As with the hierarchical mechanistic model, the GAM produces an estimate for the density of positive patients over the whole region over different times. However, unlike the hierarchical mechanistic model, it cannot explicitly provide magnitude and dispersion measures for the resistance hotspots.

### The hierarchical mechanistic Bayesian model

The model comprises three levels. First, the data level which states that the data is depends on the sampling probability and the density of positive patients. Second, the process level which captures the spatiotemporal mechanisms of the density of positive patients. Third, the parameter level which estimates the model parameters.

#### Data level

The data is an aggregate of *M* studies, which each have a unique location and time. The *M* studies are indexed by *i* = 1, 2, …, *M*. We represented the data as *y*_*i*_ ∈ ***Z***, which is the number of positive patients in study *i*. We model this as

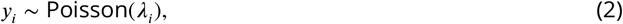

where *λ*_*i*_ > 0 is a latent spatiotemporal process defined by

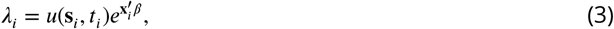

where *u*(**s**_*i, ti*_) is the density of positive patients, at the location and time of study *i* (determined by the Process level). The total record of observed counts of positive patients is ***Y***. A patient at the location and time of the study *i* has a probability of being sampled. This probability depends on patient covariates, such as age, which are captured by **x**_*i*_. The *β* in Eq. 3 are the regression coefficients (determined by the Parameter level) for the individual covariates **x**_*i*_. Therefore, the data level, Eq. 2 and Eq. 3, states that the data *y*_*i*_ depends on the density of positive patients, and a probability of being sampled.

#### Process level

The density of positive patients is a dynamic process captured by the partial differential equation (PDE)

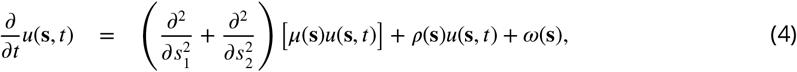

where the diffusion rate log(*μ*(**s**)) = *α*_0_ + **a**(**s**)′*α* depends on spatial covariates captured by **a**(**s**), and the growth rate *ρ*(**s**) = *γ*_0_ + **c**(**s**)′*γ* depends on spatial covariates captured by **c**(**s**). The *α*_0_, *α, γ*_0_, and *γ* are the regression coefficients (determined by the Parameter level) for the spatial covariates **a**(**s**) and **c**(**s**). The first two terms of Eq. 4, the diffusion and growth terms, make up the ecological diffusion equation, which is often used to model animal movement using abundance data ***Garlick et al. (2011)***; ***Hooten et al. (2013)***; ***Hooten and Wikle (2010)***.

For our application here, we no longer use the terminology ‘diffusion’ and ‘growth’ rate because they are misleading. In terms of the spread of drug resistant pathogens, *μ*(**s**) relates to transmission to neighbouring regions and the *ρ*(**s**) relates to transmission within a local area, see Fig. 2. With regards to the choice of spatial covariates, the transmission of drug resistant pathogens depends on the prevalence of the disease, thus **a**(**s**) and **c**(**s**) include the disease prevalence.

**Figure 2.**
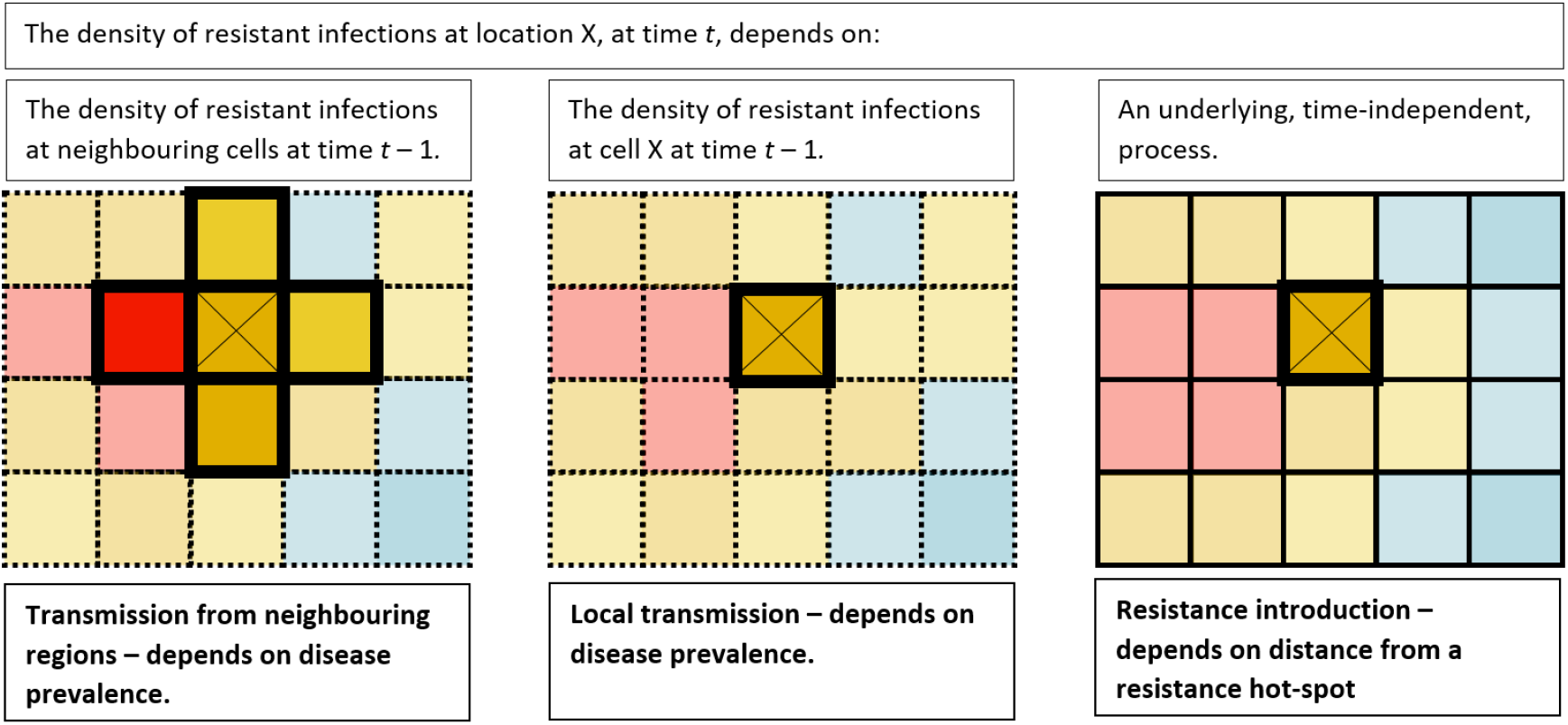
Visual description of each of three terms in Eq. 4 in relation to the spread of resistant pathogens. The highlighted boxes indicate the locations that affect the density at location X.

The last term of Eq. 4, *ω*(**s**), is an additional term we added to account for the underlying time-independent component which assumes mutations associated with resistance originates from hotspots, see Eq. 1. We use zero boundary conditions, and initial conditions *u*(**s**, 0) = *ω*(**s**), meaning that initially, the only influence on the emergence of drug resistant pathogens is the distance from a hotspot.

#### Parameter level

To complete the Bayesian specification of the spatiotemporal model, we describe the probability models for the parameters discussed in the data and process levels. The parameters which require prior distributions include the magnitude and dispersal of the resistance hotspots, ***θ*** and ***ϕ***, which we assign priors *θ*_*n*_ ∼ TN(0, 10^6^) and *ϕ*_*n*_ ∼ TN(0, 10^6^) where *n* ∈ [1, *N*] and TN refers to a normal distribution truncated below zero).

For regression coefficients ***β, α*** and ***γ*** we used priors drawn from a normal distribution with mean 0 and variance 10, as in ***Hefley et al. (2017)***: ***β*** ∼ *N*(0, 10**I**), *α*_0_ ∼ *N*(0, 10), *α* ∼ *N*(0, 10**I**), *γ*_0_ ∼ *N*(0, 10), and *γ* ∼ *N*(0, 10**I**), where **I** is the identity matrix.

### Numerical implementation

The parameters are determined using a Monte Carlo Markov Chain (MCMC), coded in R. From these parameter values, we can recover the spatiotemporal density of positive patients. The model algorithm is provided below, followed by details about the creation of the simulated data we used to demonstrate the model application. All codes are provided on GitHub. They are built upon the codes provided by ***Hefley et al. (2017)***. We run the model for 250,000 iterations and remove the first 1,000 iterations due to burn-in.

#### The algorithm

For each MCMC iteration, the estimates for the parameters are updated using Metropolis Hastings. The algorithm is,

1. Set initial values for ***α***, *β*, ***γ***, ***θ, ϕ***
2. **while** *l* < *m* **do**
3. update *u*(**s**, *t*)
4. sample [***θ, ϕ*** |***α***, *β*, ***γ***]
5. update *u*(**s**, *t*)
6. sample [***α*** | *β*, ***γ***, ***θ, ϕ***]
7. update *u*(**s**, *t*)
8. sample [*β* | ***α, γ***, ***θ, ϕ***]
9. sample [***γ*** | ***α***, *β*, ***θ, ϕ***]
10. **end while**

Updating *u*(**s**, *t*) requires solving a PDE (Eq. 4) over a fine-scale grid, which is very computationally expensive. We apply the homogenisation technique ***Garlick et al. (2011)***; ***Powell and Zimmermann (2004)***; ***Hooten et al. (2013)*** which means that each time *u*(**s**, *t*) is updated (steps 3, 5, and 7 above), the process is

a. Calculate *μ*(**s**), *ρ*(**s**) and 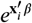 using current estimates for the regression coefficients.
b. Convert *μ*(**s**) and *ρ*(**s**) to a coarser grid.
c. Solve the PDE (Eq. 4) on the coarser grid.
d. Convert the solution on the coarse grid to the original fine-scale grid.

#### Creating the simulated data

To demonstrate the utility of the model we created simulated data over a unit square with five resistance hotspots. We assume that the hotspot locations are known, but this is not a restriction of the model. The locations are *d*_1_ = (0.1, 0.9), *d*_2_ = (0.1, 0.1), *d*_3_ = (0.5, 0.5), *d*_4_ = (0.6, 0.2) and *d*_5_ = (0.9, 0.5).

We set the magnitudes of the hotspots to *θ*_1_ = 80, *θ*_2_ = 70, *θ*_3_ = 65, *θ*_4_ = 60 and *θ*_5_ = 60. We set the dispersal of each hotspot to *ϕ*_1_ = 0.08, *ϕ*_2_ = 0.09, *ϕ*_3_ = 0.1, *ϕ*_4_ = 0.15 and *ϕ*_5_ = 0.12. These values were chosen to include an example where the hotspot with the greatest magnitude (hotspot 1) is isolated so resistance does not disperse far, see Fig. 1. In our demonstration, the covariates **a**(**s**) and **c**(**s**) are each a single covariate that varies in space. When estimating drug resistance, the main covariate should be the prevalence of the disease which has a non-random spatial pattern. To simply represent this, both **a**(**s**) and **c**(**s**) are set to values between −0.5 and 0.5 which are ordered so that the greater values are at the bottom of the square, graduating towards the top of square, see Fig. 3. Although we generated **a**(**s**) and **c**(**s**) in exactly the same manner for our demonstration, we continue with distinct notation for clarity, and to serve as a reminder that the model allows for two distinct spatial covariates which affect neighbouring and local transmission differently. The transmission to neighbouring regions *μ*(**s**) depends on the spatial covariate **a**(**s**),

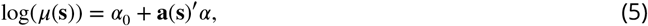

where we set the coefficients to *α*_0_ = −8 and *α*_1_ = 1. The local transmission *ρ*(**s**) depends on the spatial covariate **c**(**s**),

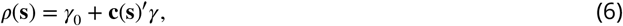

where we set the coefficients to *γ*_0_ = 0.2 and *γ*_1_ = 0.1. The simulated density of positive patients for 20 years, using Eq. 4, gives Fig. 4. Notice that although hotspot 1 (the top left hotspot) has the highest magnitude, the region surrounding hotspot 3 (the middle hotspot) has the highest density of positive patients. This demonstrates that simply observing the presence of positive patients can mask underlying spatial dynamics.

**Figure 3.**
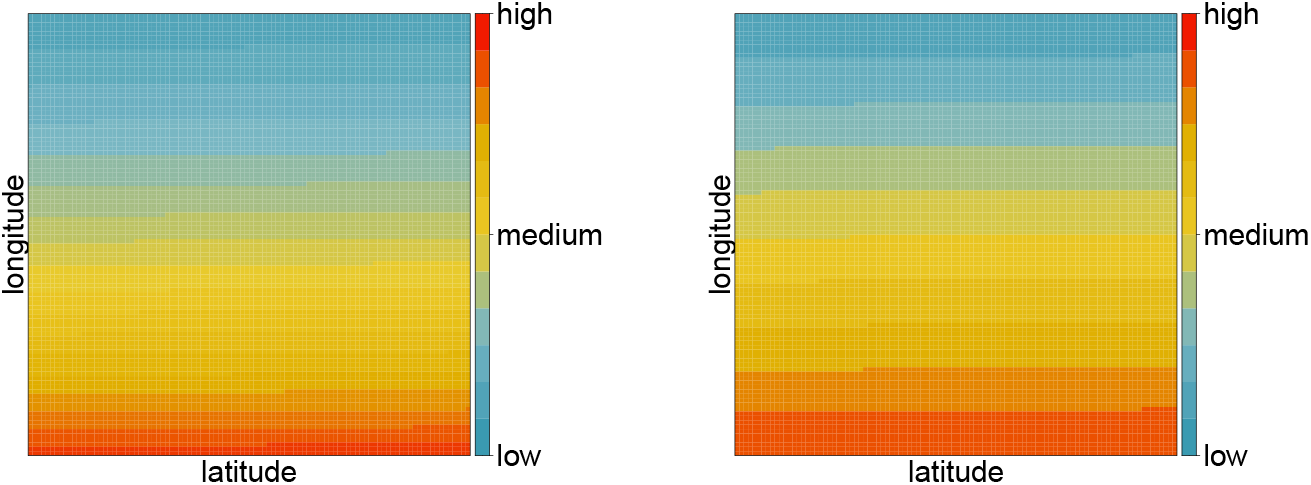
The transmission of drug resistant pathogens depends on spatial covariates **a**(**s**) and **c**(**s**), which represent covariates such as disease prevalence. Left: The transmission to neighbouring regions, *μ*(**s**), where log(*μ*(**s**)) = *α*_0_ + **a**(**s**)′*α*. Right: The local transmission, *ρ*(**s**), where *ρ*(**s**) = *γ*_0_ + **c**(**s**)′*γ*. Since this is a demonstration, the pattern is relevant (transmission of drug resistant pathogens is easier in the south than in the north), but the actual values are not, so they are omitted for clarity.

**Figure 4.**
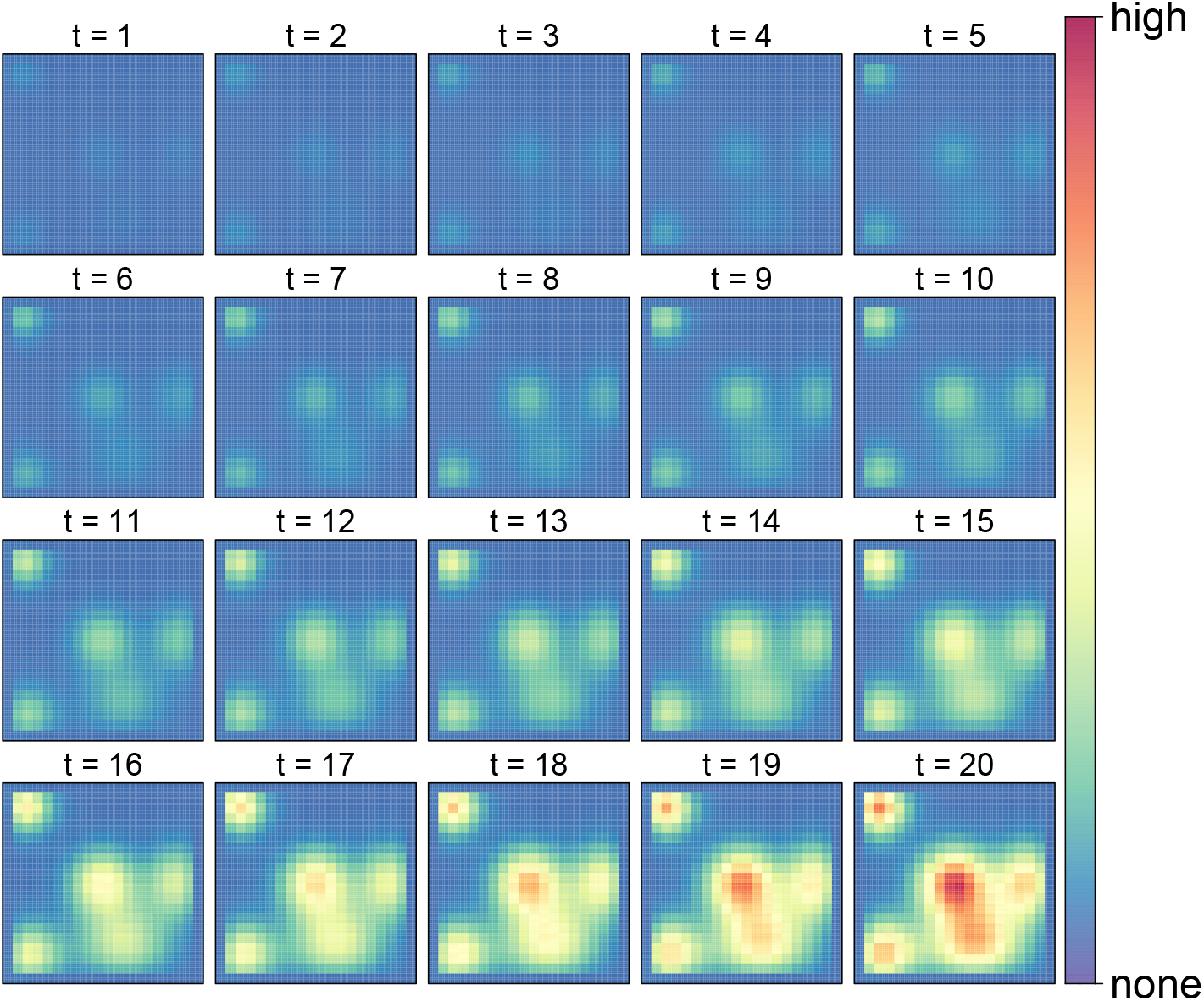
The simulated density of drug resistant infections. The values are not relevant for this simulated example so they are omitted for clarity.

The probability that a positive patient is sampled is given by

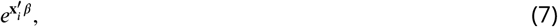

where **x**_*i*_ contains patient information, such as the patient age. In our demonstration, **x**_*i*_ is a single covariate containing random values between −0.5 and 0.5, and *β* = −10 (note that there is not an intercept term in **x**_*i*_). With biased sampling, as is the current approach, we set the sampling probability to be one where resistance is present, and random otherwise, see the top plot of Fig. 5. With unbiased sampling, each location is equally likely to be sampled, see the bottom plot of Fig. 5. We assume that the same proportion of the region is sampled each year, either 5%, 10%, 15% 20% and 30%, however this is not a restriction of the model.

**Figure 5.**
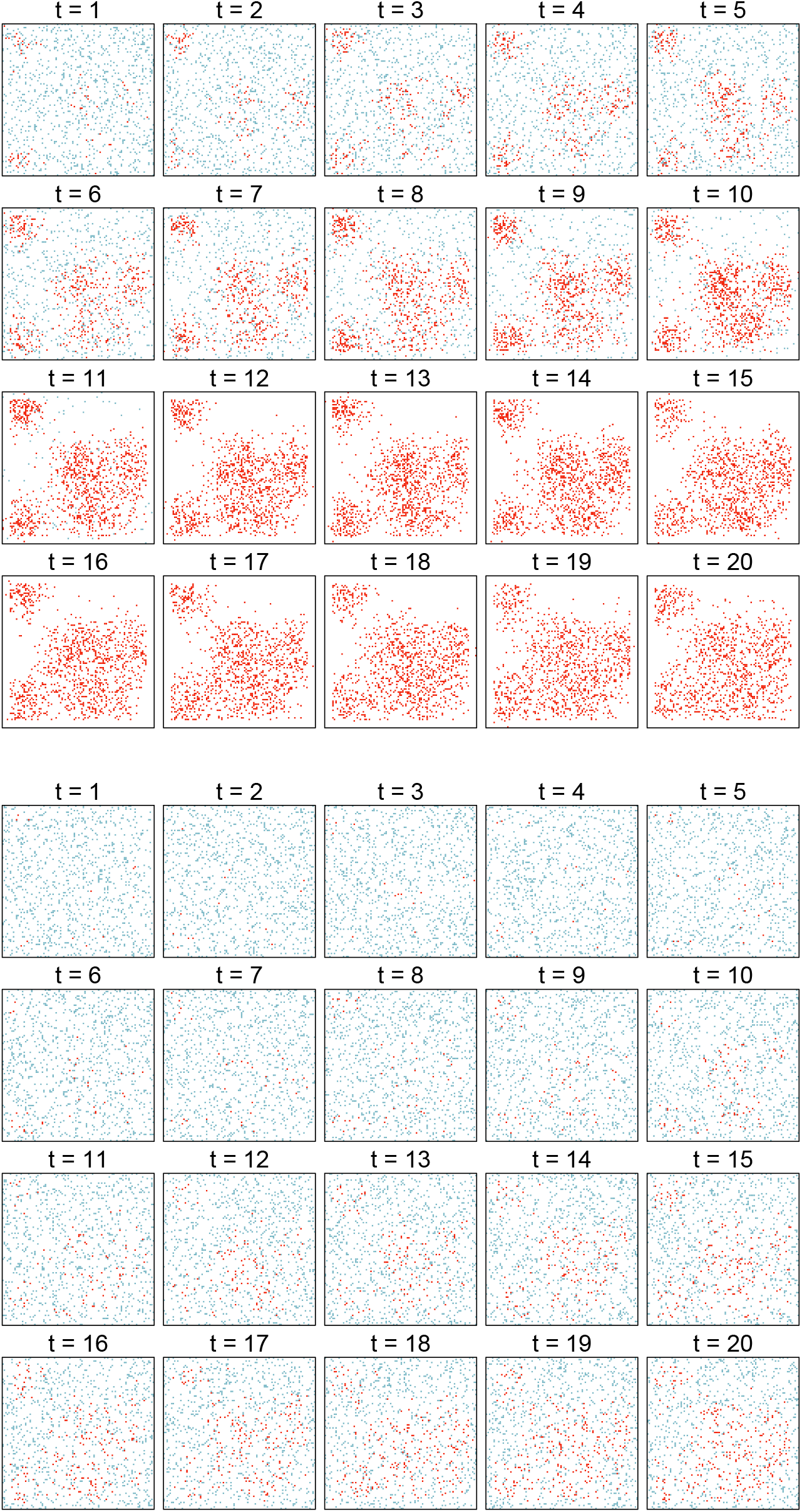
Bias and unbias sampling where 10% of region is sampling. For biased sampling, patients who are assumed to have a drug-resistant infection are more likely to be sampled. Red dots refer to a patient testing positive for a drug-resistant infection. Blue dots refer to a patient testing negative for a drug-resistant infection.

**Figure 6.**
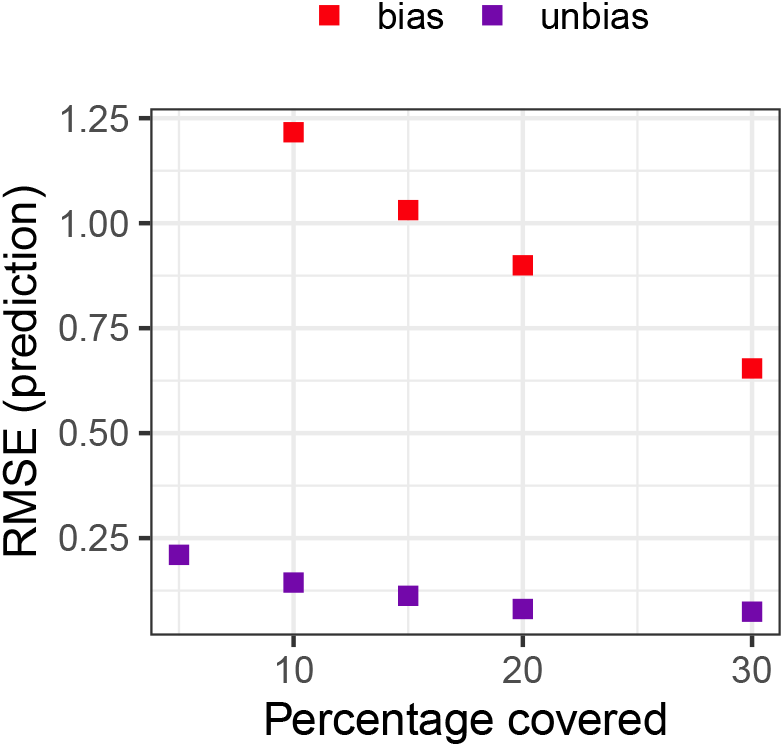
The root mean squared error (RMSE) over the 20 year period for different sampling techniques (bias, unbias), using the hierarchical mechanistic model.

**Figure 7.**
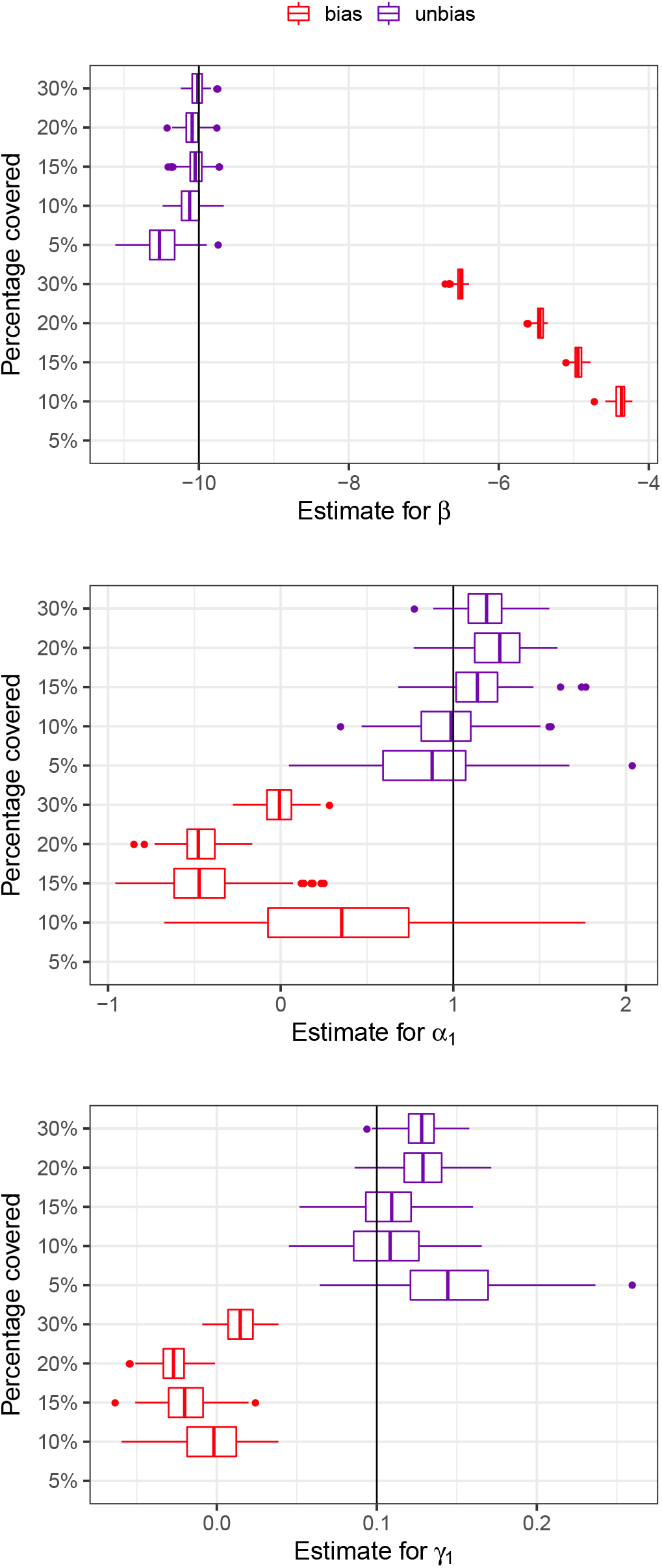
The estimates from the MCMC for the regression coefficients: *β* relating to the sampling probability, *α*_1_ relating to transmission to neighbouring regions, and *γ*_1_ relating to local transmission. The actual value is indicated by a solid line. The model fails when only sampling 5% of the region in a biased manner.

**Figure 8.**
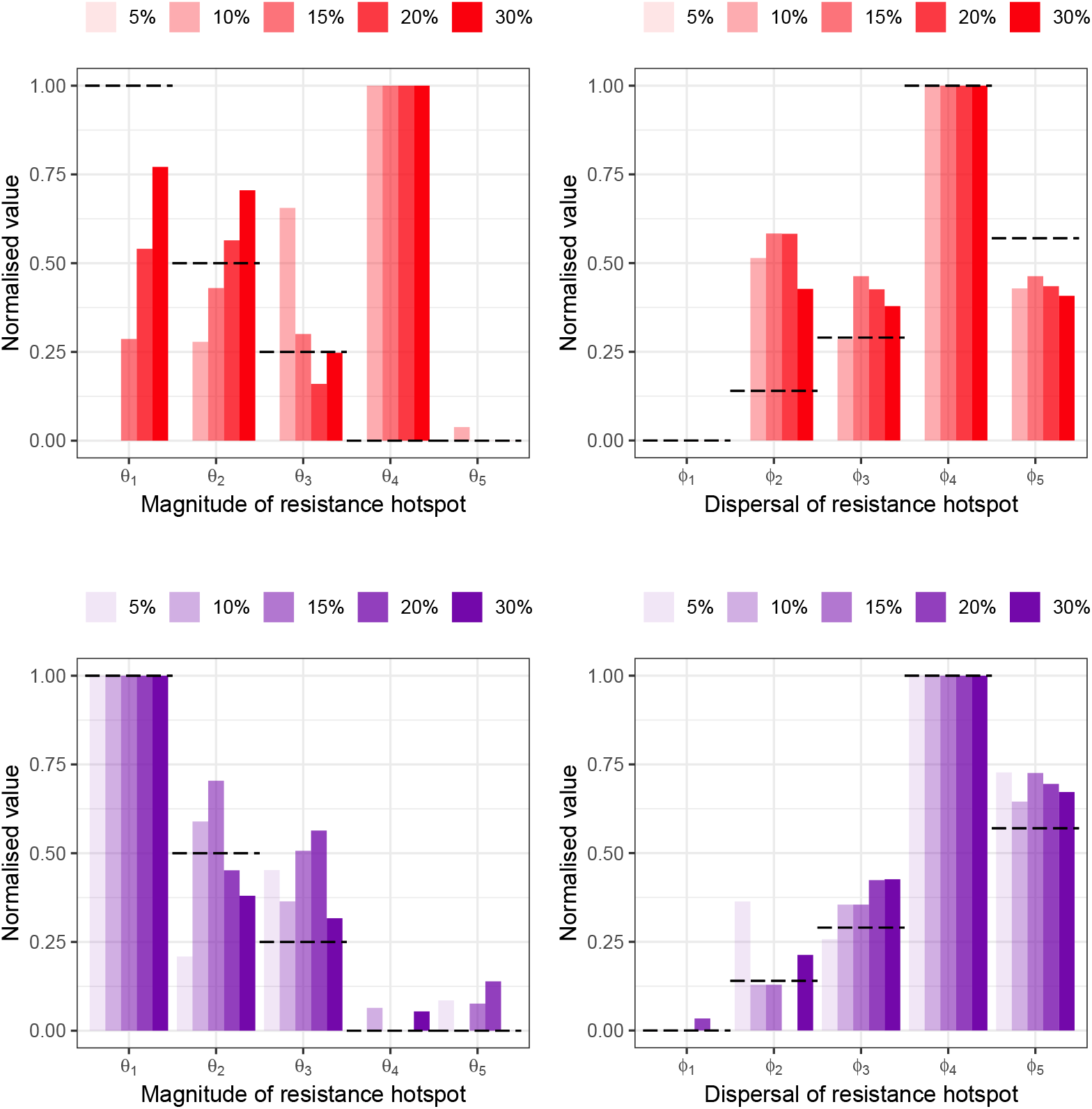
The normalised estimates from the MCMC for the magnitudes of the hotspots (left) and disperal of the hotspots (right) with biased sampling (red) and unbiased sampling (purple). The actual normalised value is indicated by the dashed line. The proportion of the region sampled is indicated by the intensity of the colour. The model fails when only sampling 5% of the region in a biased manner.

### Comparison with generalised additive models

As with the hierarchical mechanistic model, a spatiotemporal generalised additive model (GAM) produces an estimate for the whole region over different times, but it cannot explicitly provide magnitude and dispersion measures for the resistance hotspots. Our GAM assumes that the data, *y*_*i*_ ∈ ***Z*** being the number of positive patients in study *i*, is modelled such that

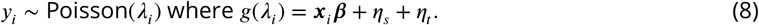

The probability that a patient is positive, *λ*_*i*_, is transformed using a link function *g*(·) and depends on both the individual data (such as the patient age), and spatial covariates (such as disease prevalence) which are included in the vector ***x***_*i*_. Note that unlike the hierarchical mechanistic model, the individual and spatial covariates are treated the same. However, we can still explicitly include the effect of time, *η*_*t*_, and spatial location, *η*_*s*_, albeit they are modelled individually and do not depend on the covariates. To illustrate the importance of explicitly including the effect of time and space, we also used a GAM that did not explicitly state these components,

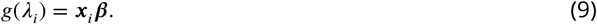

### Model assessment

To evaluate the hierarchical mechanistic model and the GAM we use a standard measure of accuracy: the root mean squared error (RMSE) of the predicted true density of positive patients in space and time. We also examine the accuracy of the models by comparing the estimated regression coefficients with the actual values used when creating the simulated data.

Of most interest for drug resistance, with the hierarchical mechanistic model, we recover the magnitudes and dispersals of the hotspots. For strategic decisions, such as checking the quality and procedures of a particular health care centre, the actual magnitude of the hotspot is irrelevant. Only the ranking of this hotspot compared to the others is required. Therefore we focus on recovering the ranking of the magnitudes of the hotspots. For completeness, we also recover the ranking of the dispersals of the hotspots.

To highlight the ranking of hotspots, and not the actual magnitude, in Results we present the normalised values: 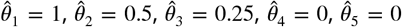, and 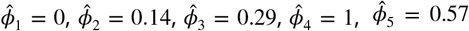, where the 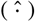denotes the normalised value. See Creating th_5_e simulated data for the original magnitude and dispersal values. We do not recover the magnitudes and dispersals of the hotspots when applying the GAM because the GAM cannot recover this information.

## Acknowledgments

This research was funded by Tamsin Lee’s Marie Curie Individual Fellowship 839121, Horizon 2020. Mevin Hooten provided early modelling guidance. Calculations were performed at sciCORE (http://scicore.unibas.ch/) scientific computing center at University of Basel.

## References

Boni MF, Smith DL, Laxminarayan R. Benefits of using multiple first-line therapies against malaria. Proceedings of the National Academy of Sciences. 2008; 105(37):14216–14221.

Brookes VJ, Hernádez-Jover M, Black PF, Ward MP. Preparedness for emerging infectious diseases: pathways from anticipation to action. Epidemiology and Infection. 2015; 143(10):2043–2058. doi: 10.1017/S095026881400315X.

Buckee CO, Wesolowski A, Eagle NN, Hansen E, Snow RW. Mobile phones and malaria: modeling human and parasite travel. Travel medicine and infectious disease. 2013; 11(1):15–22.

Conn PB, Johnson DS, Hoef JMV, Hooten MB, London JM, Boveng PL. Using spatiotemporal statistical models to estimate animal abundance and infer ecological dynamics from survey counts. Ecological Monographs. 2015; 85(2):235–252.

Davison HC, Woolhouse ME, Low JC. What is antibiotic resistance and how can we measure it? Trends in microbiology. 2000; 8(12):554–559.

Gardy JL, Loman NJ. Towards a genomics-informed, real-time, global pathogen surveillance system. Nature Reviews Genetics. 2018; 19(1):9–20.

Garlick MJ, Powell JA, Hooten MB, McFarlane LR. Homogenization of large-scale movement models in ecology. Bulletin of Mathematical Biology. 2011; 73(9):2088–2108.

Hefley TJ, Hooten MB, Russell RE, Walsh DP, Powell JA. When mechanism matters: Bayesian forecasting using models of ecological diffusion. Ecology Letters. 2017; 20(5):640–650.

Hooten MB, Garlick MJ, Powell JA. Computationally efficient statistical differential equation modeling using homogenization. Journal of agricultural, biological, and environmental statistics. 2013; 18(3):405–428.

Hooten MB, Wikle CK. A hierarchical Bayesian non-linear spatio-temporal model for the spread of invasive species with application to the Eurasian Collared-Dove. Environmental and Ecological Statistics. 2008; 15(1):59–70.

Hooten MB, Wikle CK. Statistical agent-based models for discrete spatio-temporal systems. Journal of the American Statistical Association. 2010; 105(489):236–248.

Iskandar K, Molinier L, Hallit S, et al. Surveillance of antimicrobial resistance in low- and middle-income countries: a scattered picture. Antimicrobial Resistance & Infection Control. 2021; 10:63. https://doi.org/10.1186/s13756-021-00931-w, doi: 10.1186/s13756-021-00931-w.

Knight GM, Davies NG, Colijn C, Coll F, Donker T, Gifford DR, Glover RE, Jit M, Klemm E, Lehtinen S, et al. Mathematical modelling for antibiotic resistance control policy: do we know enough? BMC infectious diseases. 2019; 19(1):1–9.

Mari A, Roloff TC, Stange M, Soegaard KK, Asllanaj E, Tauriello G, Alexander LT, Schweitzer M, Leuzinger K, Gensch A, et al. Global surveillance of potential antiviral drug resistance in SARS-CoV-2: proof of concept focussing on the RNA-dependent RNA polymerase. medRxiv. 2021; p. 2020–12.

Powell JA, Zimmermann NE. Multiscale analysis of active seed dispersal contributes to resolving Reid’s paradox. Ecology. 2004; 85(2):490–506.

Roberts SL. In: Romaniuk S, Thapa M, Marton P, editors. Emerging and Re-emerging Diseases Cham: Springer International Publishing; 2019. p. 1–9. https://doi.org/10.1007/978-3-319-74336-3_531-1, doi: 10.1007/978-3-319-74336-3_531-1.

Wikle CK. Hierarchical Bayesian models for predicting the spread of ecological processes. Ecology. 2003; 84(6):1382–1394.

Wikle CK, Hooten MB. Hierarchical Bayesian spatio-temporal models for population spread. Applications of computational statistics in the environmental sciences: hierarchical Bayes and MCMC methods. 2006; 145169.

World Health Organization. Future Surveillance for Epidemic and Pandemic Diseases: A 2023 Perspective. Geneva: World Health Organization; 2023. https://www.who.int/publications/i/item/9789240080959, licence: CC BY-NC-SA 3.0 IGO.

